# Intralysosomal pathogens differentially influence the proteolytic potential of their niche

**DOI:** 10.1101/2025.05.21.655381

**Authors:** Lauren E. Bird, Bangyan Xu, Erin N. S. McGowan, Patrice Newton, Nichollas E. Scott, Malcolm J. McConville, Laura E. Edgington-Mitchell, Hayley J. Newton

## Abstract

Avoiding lysosomal degradation is vital to the success of intracellular pathogens. The Gram-negative bacterium *Coxiella burnetii* and protozoan parasites of the *Leishmania* genus are unique in being able to replicate within the mature phagolysosomal compartment of host cells, though the exact mechanisms utilised to withstand this hostile environment are not clearly defined. We recently reported that *C. burnetii* removes the lysosomal protease cathepsin B during infection of mammalian cells. Here, we aimed to determine if this virulence strategy was also employed by the intralysosomal pathogen, *Leishmania mexicana*. In contrast to *C. burnetii*, decreases in the activity of specific cathepsins were not detected in *L. mexicana*-infected host cells as determined using immunoblotting and protease activity-based probes. Co-infection of THP-1 macrophage-like cells with both pathogens resulted in a proteolytic and secretory phenotype consistent with *C. burnetii* infection, suggesting that *C. burnetii-*induced remodelling of the lysosome is not influenced by *L. mexicana*. The host cell proteome and secretome of *L. mexicana* infected cells was defined using mass spectrometry. This confirmed that, unlike *C. burnetii*, *L. mexicana* does not induce increased abundance of lysosomal proteins either intracellularly or in the extracellular milieu. Collectively, this study reveals that although *C. burnetii* and *L. mexicana* reside in a phagolysosomal intracellular niche, they employ divergent mechanisms to survive within this hostile compartment.

## INTRODUCTION

Evading lysosomal degradation is of critical importance to successful intracellular pathogens. The lysosome, classically considered the waste disposal and recycling centre of the cell, is a hostile environment consisting of many hydrolytic enzymes which serve to break down macromolecules to maintain cellular homeostasis. Most intracellular pathogens have evolved strategies to avoid delivery to lysosomes, either by halting maturation of the pathogen-containing phagosome (e.g. *Legionella pneumophila, Mycobacterium tuberculosis*) [1] or escaping into the cytosol for replication (*Rickettsia*, *Listeria, Shigella* spp.) [2]. The only known human pathogens which replicate inside a lysosome-derived niche are the protozoa *Leishmania* spp. and the Gram-negative bacterium *Coxiella burnetii*.

*Leishmania* spp. are protozoan parasites which cause the devastating and potentially fatal spectrum of diseases collectively termed ‘leishmaniases’ (reviewed in [3]). Symptoms of leishmaniasis can range from self-healing cutaneous lesions to chronic, multisystemic disease depending on the parasite species responsible for infection. An estimated 700,000-1 million new cases occur annually, with the World Health Organization listing leishmaniases as a class of major neglected tropical diseases [4].

*Leishmania* parasites exhibit a digenetic lifecycle. The extracellular, flagellated promastigote replicates in the midgut of the sandfly and is transmitted to a vertebrate host during a bloodmeal. Promastigotes are internalised by phagocytic cells and subsequently differentiate into aflagellated, non-motile amastigotes, which replicate inside an intracellular parasitophorous vacuole (PV) with phagolysosomal features [5]. Macrophages and monocytes are the primary cell type infected by *Leishmania,* though parasites can also enter and persist in neutrophils and dendritic cells, as well as non-phagocytic cells [6, 7].

Following phagocytosis, parasites are delivered to a mature phagolysosome which continues to fuse with other vesicles of the endolysosomal system. Some *Leishmania* species (e.g. *L. amazonensis, L. mexicana*) generate large communal phagolysosomes containing hundreds of parasites, while other species (e.g. *L. major*) reside within individual, tight fitting phagolysosomes [8]. During maturation, *Leishmania* PVs sequentially acquire the endosomal markers Rab5 and Rab7, as well as lysosomal membrane proteins (LAMP-1, LAMP-2) and hydrolases (cathepsins) [9, 10, 11, 12]. The promastigote stages of some species of *Leishmania* may transiently delay the maturation of the PV, although this is not evident with *L. mexicana* promastigotes or the amastigote stages of all species (reviewed in [13]). How *Leishmania* amastigotes to avoid hydrolytic degradation in a lysosome-derived compartment remains poorly understood.

The only other known pathogen which requires a lysosomal environment for replication is the Gram-negative bacterium *C. burnetii*. *C. burnetii* is the causative agent of Q fever, which has a spectrum of clinical presentations ranging from asymptomatic seroconversion to chronic, life-threatening illness [14]. Human infection occurs through inhalation of contaminated aerosols, and the bacterium establishes its replicative niche inside alveolar macrophages of infected individuals. The mature pathogen-containing compartment is termed the *Coxiella-* containing vacuole (CCV) and displays both endolysosomal (Rab5, Rab7, LAMP-1 and 2) and autophagosomal (LC3, p62) markers [15]. From within this replicative niche, *C. burnetii* employs a type IV-B secretion system (T4SS) to translocate approximately 130 bacterial proteins into the host cytosol to manipulate various host processes which contribute to infection [16].

The *C. burnetii* CCV demonstrates remarkable fusogenicity and has been previously shown to take up inert latex beads or other pathogens including *Trypanosoma cruzi* [17] and *Mycobacterium avium* [18]. There is also existing evidence that *C. burnetii* and *L. amazonensis* can cohabit a single vacuole within a mammalian cell. It has been demonstrated that *L. amazonensis* can traffic to and replicate within pre-existing *C. burnetii* vacuoles in Chinese Hamster Ovary cells [19], and that Vero cells infected with *L. amazonensis* can be super-infected with Dot/Icm-deficient *C. burnetii* to promote intracellular proliferation of this normally non-replicative strain [20]. This suggests that the vacuoles housing *C. burnetii* and *L. amazonensis* are at least superficially similar. Whether this is true of other *Leishmania* species is unclear.

We recently reported that *C. burnetii* removes the lysosomal protease cathepsin B from infected cells, leading to improved bacterial replication and intracellular success [21]. In this study, we have investigated whether *L. mexicana* similarly manipulate the levels of specific lysosomal proteases. Using a panel of activity-based probes (ABPs) and immunoblotting, it was found that *C. burnetii* and *L. mexicana* differentially modulate the proteolytic capacity of their replicative niche. Additionally, *L. mexicana* infection does not induce the extracellular secretion of lysosomal content, as previously observed for *C. burnetii* [21]. Collectively, this indicates that although these pathogens reside inside a niche with superficially similar characteristics, they modulate the cohort of lysosomal proteases in distinct ways to promote their own intracellular survival.

## RESULTS

Cathepsin abundance and activity is differentially modulated by *C. burnetii* and *L. mexicana C. burnetii* elicits removal of the lysosomal protease cathepsin B from mammalian cells to promote infection [21]. To explore whether this is a shared strategy of intralysosomal pathogens, the abundance of specific cathepsins was measured during infection with *L. mexicana*. *L. mexicana* was chosen as a representative leishmanial species because of the similar fusogenicity of the *L. mexicana* PV to the *C. burnetii* CCV (multiple parasites within a single communal vacuole) and the ability to work with this pathogen under Biosafety Level 2 (BSL-2) conditions. Differentiated THP-1 cells were infected with either *C. burnetii* or *L. mexicana* at a multiplicity of infection (MOI) of 10 and the abundance of specific cathepsins was assessed over 5 days of infection. Consistent with previous observations, cathepsin B was undetectable in *C. burnetii-*infected cells (Fig 1A, B). In contrast, THP-1 cells infected with *L. mexicana* exhibited an increase in cathepsin B abundance over the infection time-course (Fig 1A, B). To determine if this was the case for other cathepsins, we immunoblotted for cathepsins C and D. Consistent with previous findings [21], *C. burnetii* infection led to a reduction in cathepsin C (Fig 1A, C), while *L. mexicana* infection resulted in somewhat increased cathepsin C abundance (Fig 1A, C). The pro-form of cathepsin D (pCTSD) accumulated in *C. burnetii-*infected cells, but not uninfected or *L. mexicana*-infected cells (Fig 1A), while mature cathepsin D (mCTSD) gradually accumulated over time in both infection conditions (Fig 1D).

**Figure 1.**
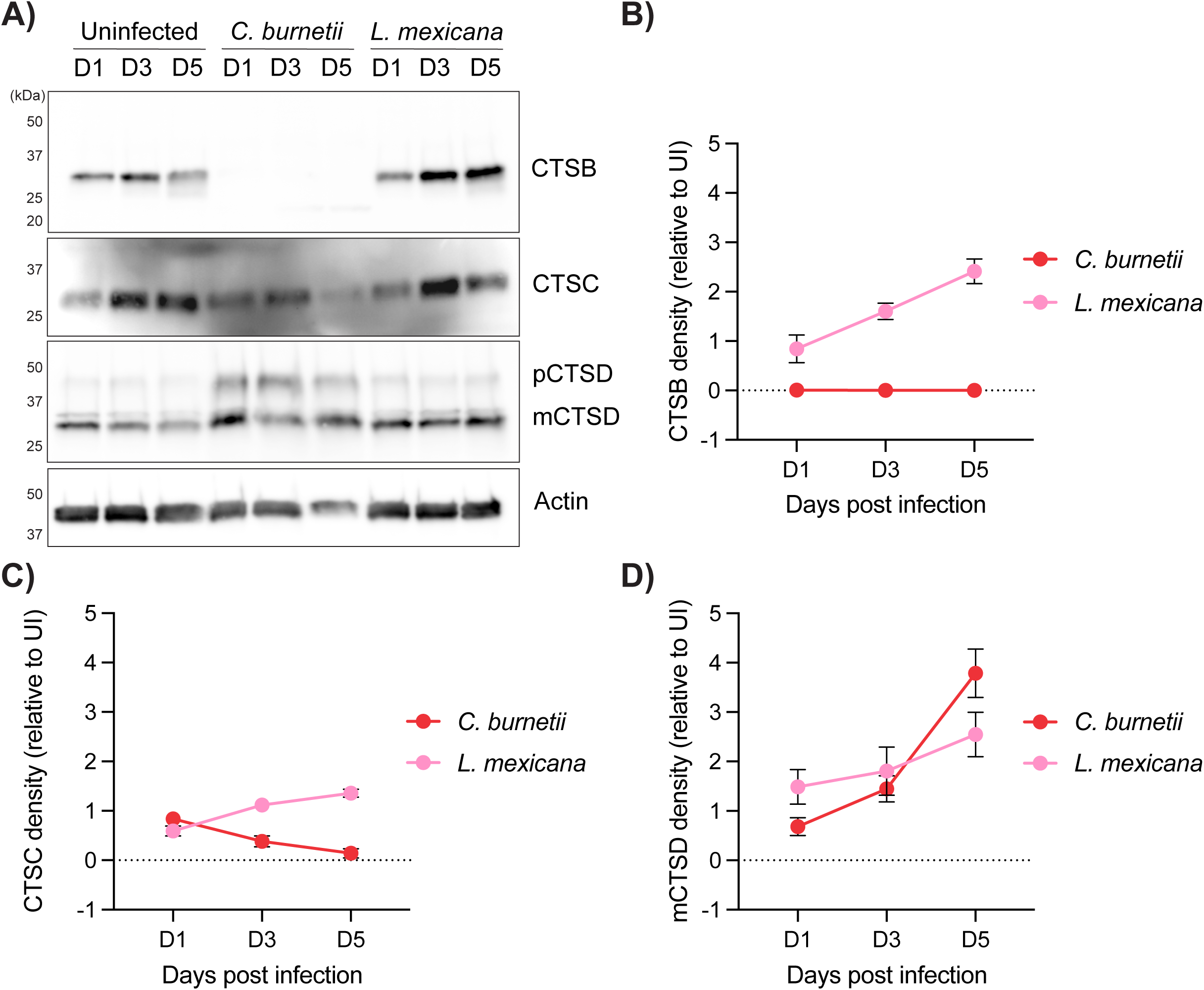
Cathepsin abundance is differentially modified by *C. burnetii* and *L. mexicana*. **(A)** THP-1 cells were left uninfected or infected for 5 days with *C. burnetii* or *L. mexicana* at a multiplicity of infection (MOI) of 10. At days 1, 3, or 5 post infection, cells were lysed and subject to SDS-PAGE and immunoblotting using antibodies against cathepsin B (CTSB), cathepsin C (CTSC), cathepsin D (CTSD) or actin as a loading control. pCTSD = pro-cathepsin D, mCTSD = mature cathepsin D. Blot is representative of 3 independent replicates. **(B)** Quantification of cathepsin B density as presented in (A). Values are expressed relative to uninfected cells at the same time point and standardised against respective actin expression. Graph depicts the average of 3 biological replicates. Error bars represent standard deviation. **(C)** Quantification of cathepsin C density as presented in (A). Values are expressed relative to uninfected cells at the same timepoint and standardised against respective actin expression. Graph depicts the average of 3 biological replicates. Error bars represent standard deviation. **(D)** Quantification of mature cathepsin D density as presented in (A). Values are expressed relative to uninfected cells at the same timepoint and standardised against respective actin expression. Graph depicts the average of 3 biological replicates. Error bars represent standard deviation.

Given that protein abundance does not necessarily correlate with enzymatic activity, these investigations were extended to examine cathepsin activity in infected cells by leveraging the activity-based probe (ABP) BMV109 [22, 23]. ABPs are small molecules which covalently bind to the active site of proteases and allow detection of enzyme activity through in-gel fluorescence [24]. BMV109 is a pan-reactive probe for cysteine cathepsin activity, allowing simultaneous detection of cathepsin B, S, X and L activity in a single sample [22, 23]. Incubation of *C. burnetii*-infected cell lysates with BMV109 again confirmed the complete loss of cathepsin B activity, consistent with its reduced abundance from cells (Fig 2A). Other cysteine cathepsins appeared broadly unchanged during *C. burnetii* infection. Furthermore, no labelling was observed when this probe was incubated with lysed *C. burnetii* alone (Fig 2B), confirming that BMV109 is specifically labelling mammalian and not *C. burnetii*-derived proteases. Interestingly, when BMV109 was applied to lysates from *L. mexicana-* infected cells, increased activity of a protease at a similar size to cathepsin S was observed (Fig 2C, “CTSS*”). This activity is likely to be associated with the well characterised cysteine proteases in *L. mexicana* amastigotes {Mottram, 2004 #457}, which was confirmed by incubating lysed *L. mexicana* amastigotes (no host cells) with the probe (Fig 2D). The other cysteine cathepsins identified with BMV109 labelling showed unchanged activity during *L. mexicana* infection (Fig 2C). Collectively, the data presented in Fig 1 and Fig 2 suggest that *C. burnetii* and *L. mexicana* differentially regulate the levels of host lysosomal proteases, despite residing within intracellular vacuoles with similar characteristics.

**Figure 2.**
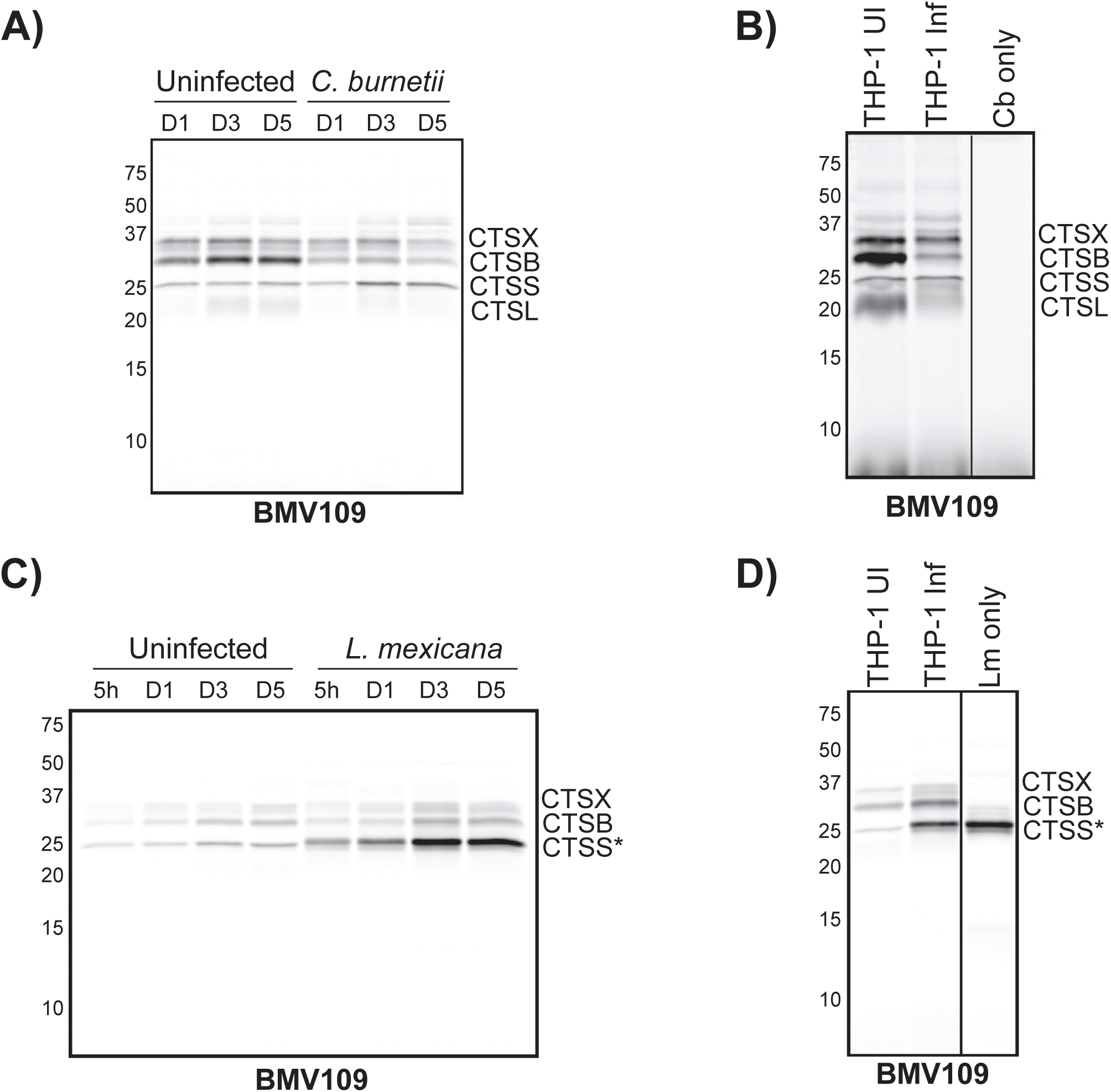
Activity-based probe profiling of cathepsin proteases during *C. burnetii* or *L. mexicana* infection of THP-1 cells. **(A)** THP-1 cells were uninfected or infected with *C. burnetii* for 5 days at a MOI of 10. At days 1, 3 or 5 post infection, cells were lysed in a citrate buffer and active cysteine proteases labelled with 1 μM BMV109 before being resolved by SDS-PAGE. Image depicts in-gel fluorescence of BMV109-labelled cysteine proteases from uninfected or *C. burnetii*-infected THP-1 cells at days 1, 3 or 5 post infection. **(B)** In-gel fluorescence of BMV109-labelled cysteine proteases from uninfected THP-1 cells (“THP-1 UI”), 3 day-*C. burnetii-*infected cells (“THP-1 Inf”), or lysed *C. burnetii* (“Cb only”). **(C)** As for (A) but cells were infected with *L. mexicana* amastigotes at a MOI of 10. Lysates were harvested at 5 h, 1-, 3-, or 5-days post infection. “*” denotes parasitic protease at the same molecular weight as CTSS. **(D)** In-gel fluorescence of BMV109-labelled cysteine proteases from uninfected THP-1 cells (“THP-1 UI”), 3 day-*L. mexicana*-infected cells (“THP-1 Inf”), or lysed *L. mexicana* amastigotes (“Lm only”). * denotes parasitic protease at the same molecular weight as CTSS. CTSX = cathepsin X, CTSB = cathepsin B, CTSS = cathepsin S, CTSL = cathepsin L.

### Co-infection with *C. burnetii* and *L. mexicana* does not alter replication dynamics or protease abundance

We next aimed to establish a co-infection model whereby cells were simultaneously infected with *C. burnetii* (either wild-type or a T4SS-deficient mutant, *icmL*::Tn) and *L. mexicana* (Fig 3A). In our previous work, *C. burnetii* replication was negatively impacted by overexpression of cathepsin B [21]. Since cathepsin B is retained during *L. mexicana* infection (Fig 1), we speculated that co-infection might similarly affect *C. burnetii* replication.

**Figure 3.**
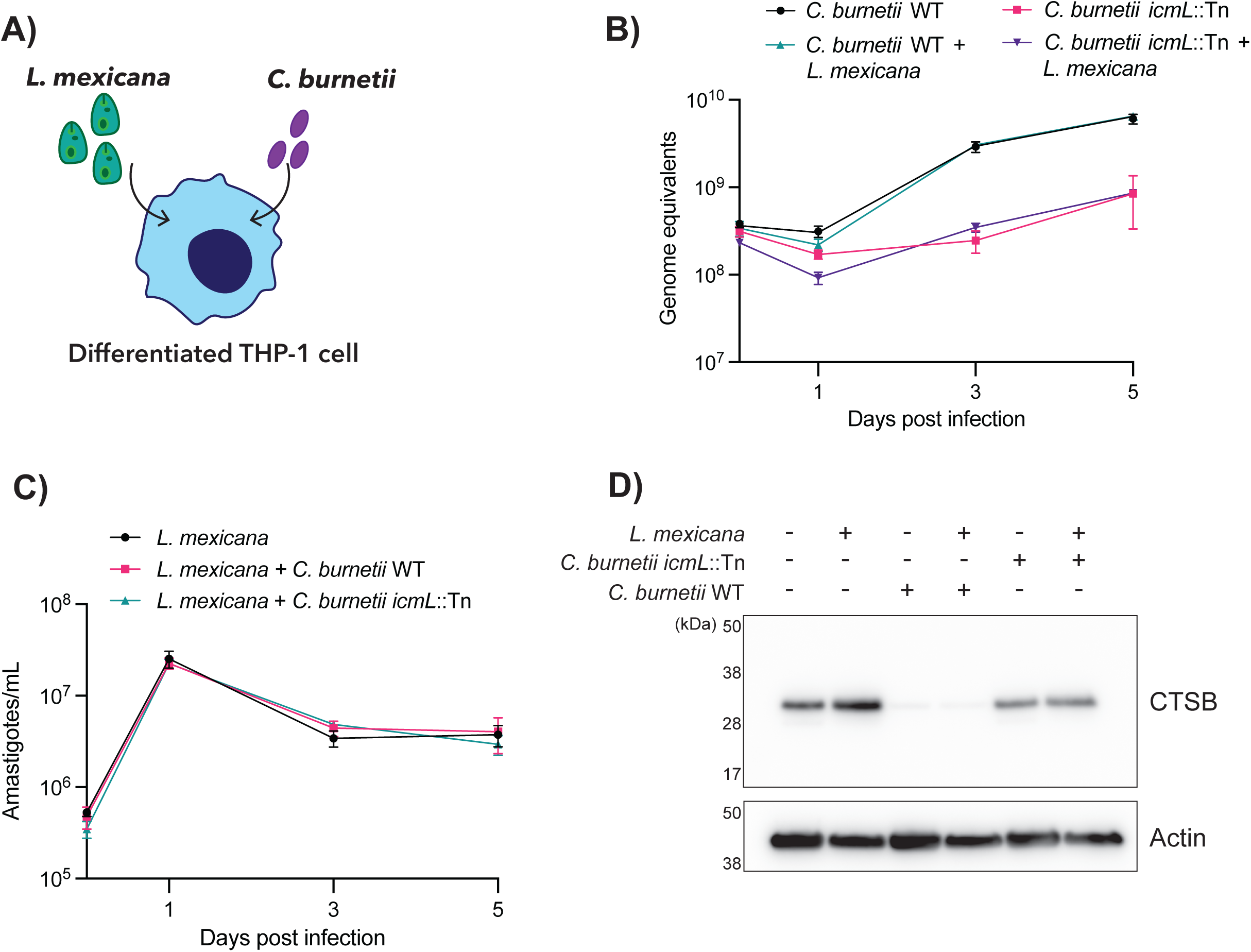
Co-infection with *C. burnetii* and *L. mexicana* does not impact replication dynamics or protease abundance. **(A)** Schematic of experimental design. **(B)** qPCR analysis of *C. burnetii* intracellular replication in THP-1 cells infected with the indicated strains +/-*L. mexicana*. Genome equivalents were calculated based on a standard curve. Individual data points represent the mean of 3 biological replicates. Error bars denote standard deviation. **(C)** qPCR analysis of *L. mexicana* intracellular replication in THP-1 cells infected +/-*C. burnetii* (wild-type or *icmL*::Tn strains). Individual data points represent the mean of 3 biological replicates. Error bars denote standard deviation. **(D)** THP-1 cells were uninfected or infected with the indicated pathogen(s) at a MOI of 10. At 3 days post infection, lysates were harvested and subjected to SDS-PAGE and immunoblotting. CTSB = cathepsin B.

To assess replication in co-infected cells, real-time quantitative PCR was performed using primers specific to the *C. burnetii* constitutively expressed *ompA* gene, as described previously [25]. Using this approach, no change to *C. burnetii* replication dynamics was observed, regardless of *L. mexicana* co-infection (Fig 3B). The same approach was used to quantify intracellular replication of *L. mexicana,* using primers targeting the kinetoplast DNA minicircle [26]. A sharp increase in parasite load was observed between day 0 and day 1 post-infection, followed by a slight drop and plateau for the remainder of the time-course (Fig 3C). Co-infection with either strain of *C. burnetii* did not significantly alter these dynamics.

Samples from day 3 of these infections were immunoblotted for cathepsin B. As before, a dramatic decrease in cathepsin B abundance was observed when cells were infected with *C. burnetii*, and this was not restored when *L. mexicana* was present. Infection with a *C. burnetii* T4SS mutant (*icmL*::Tn) led to cathepsin B retention in both singly infected and co-infected conditions. Taken together, these data indicate that *C. burnetii* maintains the ability to remove cathepsin B and replicate to equivalent numbers during co-infection as when infected alone.

### A *C. burnetii* T4SS mutant is able to replicate in *L. mexicana-*infected cells

No change to the replication dynamics of *C. burnetii* was observed when cells were simultaneously infected with both microbes. However, we wondered if this might be altered by sequential infection. It has previously been reported that *C. burnetii* can traffic to pre-existing vacuoles established by *L. amazonensis* and that this restored the replication defect observed in a T4SS-deficient *C. burnetii* strain [20]. However, the extent to which *C. burnetii* T4SS mutants can replicate inside a *Leishmania* vacuole has never been quantified. Given this, we established a super-infection model, where THP-1 cells were infected for 3 days with *L. mexicana* before being super-infected with *C. burnetii* (wild-type or *icmL*::Tn) (Fig 4A).

**Figure 4.**
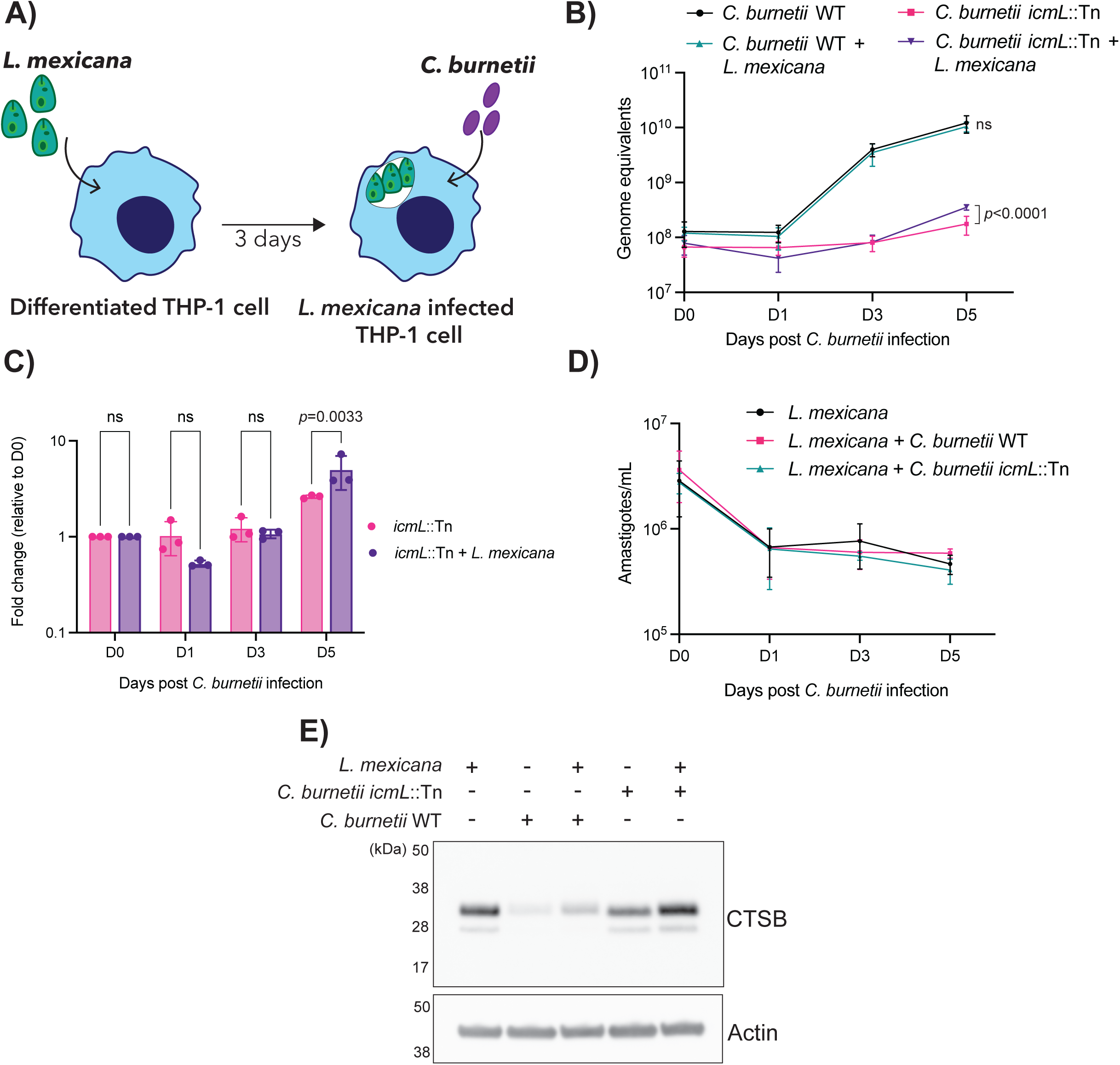
T4SS-deficient *C. burnetii* exhibit increased replication in *L. mexicana-*infected cells. **(A)** Schematic of experimental design. **(B)** qPCR analysis of *C. burnetii* intracellular replication in THP-1 cells infected with the indicated strains +/-*L. mexicana*. Genome equivalents were calculated based on a standard curve. Individual data points represent the mean of 3 biological replicates. Error bars denote standard deviation. **(C)** Data from (B) but presented as fold change in *C. burnetii* replication relative to genome equivalents present at day 0. **(D)** qPCR analysis of *L. mexicana* intracellular replication in THP-1 cells infected +/-*C. burnetii* (wild-type or *icmL*::Tn strains). Individual data points represent the mean of 3 biological replicates. Error bars denote standard deviation. **(E)** THP-1 cells were infected with *L. mexicana* for 3 days and then super-infected with *C. burnetii* for a further 3 days. At this time, lysates were harvested and subjected to SDS-PAGE and immunoblotting. CTSB = cathepsin B.

Intracellular replication of both pathogens was monitored as outlined above. The presence of a pre-existing *L. mexicana* vacuole did not alter the replication kinetics of wild-type *C. burnetii*, but did lead to a significant increase in replication of *icmL*::Tn at 5 days post *C. burnetii* infection (Fig 4B). When quantified as fold change in replication relative to bacterial load at day 0, we observed a significant increase in *C. burnetii icmL*::Tn replication at day 5 post infection when *L. mexicana* was also present (mean fold change 5.021±1.947) compared to when cells were infected with *icmL*::Tn alone (mean fold change 2.620±0.100; 2way ANOVA with Sidak’s correction, *p*=0.0033; Fig 4C). Quantification of *L. mexicana* across this time course revealed a decrease in parasite load at day 1 post *C. burnetii* infection (day 4 post *L. mexicana* infection), followed by a plateau in replication (Fig 4D).

Examination of cathepsin B abundance was performed at 3 days following *C. burnetii* infection (5 days following *L. mexicana* infection). Similar to what was observed during co-infection, cathepsin B levels were decreased when super-infected THP-1 cells were infected with wild-type *C. burnetii*, but not with *icmL*::Tn (Fig 4E). The presence of a pre-existing *L. mexicana* infection did not noticeably alter this. Altogether, these data indicate that a non-replicative T4SS mutant exhibits a small but significant increase in replication in a pre-established *L. mexicana* vacuole, though this is not due to changes in cathepsin abundance.

1. *L. mexicana* does not induce secretion of lysosomal content
2. *C. burnetii* infection causes a T4SS-independent secretion of lysosomal content into the extracellular milieu [21]. To determine if a similar phenotype was observed upon *L. mexicana* infection, proteomics analysis was performed on both the lysate and conditioned media collected from THP-1 cells after 3 days of *L. mexicana* infection (Supplementary Data 1). Using data-dependent acquisition label-free proteomics, 4,266 proteins were identified and quantified with a false discovery rate (FDR) of 1% in the whole cell lysate samples. Of these, 4,098 were host-derived and 168 were derived from *L. mexicana*. Principal component analysis showed that biological replicates from each condition clustered together (Fig 5A). Only a small number of changes to the host proteome were observed during infection, with 64 proteins significantly increased in abundance and 23 proteins showing decreased abundance, as determined by a student’s *t*-test with *p*<0.05 and log_2_ fold change>1 (Fig 5B).
3. *C. burnetii* infection leads to increased abundance of lysosomal proteins, likely as a result of nuclear translocation of transcription factors EB and E3 [27]. To determine if this also occurred during infection with *L. mexicana*, the proteomics dataset was overlaid with the Gene Ontology (GO) terms, “lysosome”, “lysosome lumen” or “lysosome membrane” (GO:0005764, GO:0043202 or GO:0005765, respectively). Unlike what has been reported for *C. burnetii* infection, there was no increased abundance of lysosomal proteins in cells infected with *L. mexicana* (Fig 5B, pink circles). Specific examination of cathepsin abundance also revealed that cathepsins were not significantly altered during *L. mexicana* infection, consistent with our earlier findings (Fig 5B, labels). This suggests that while *L. mexicana* resides in a phagolysosomal niche, there is likely no associated increase in lysosomal biogenesis.

**Figure 5.**
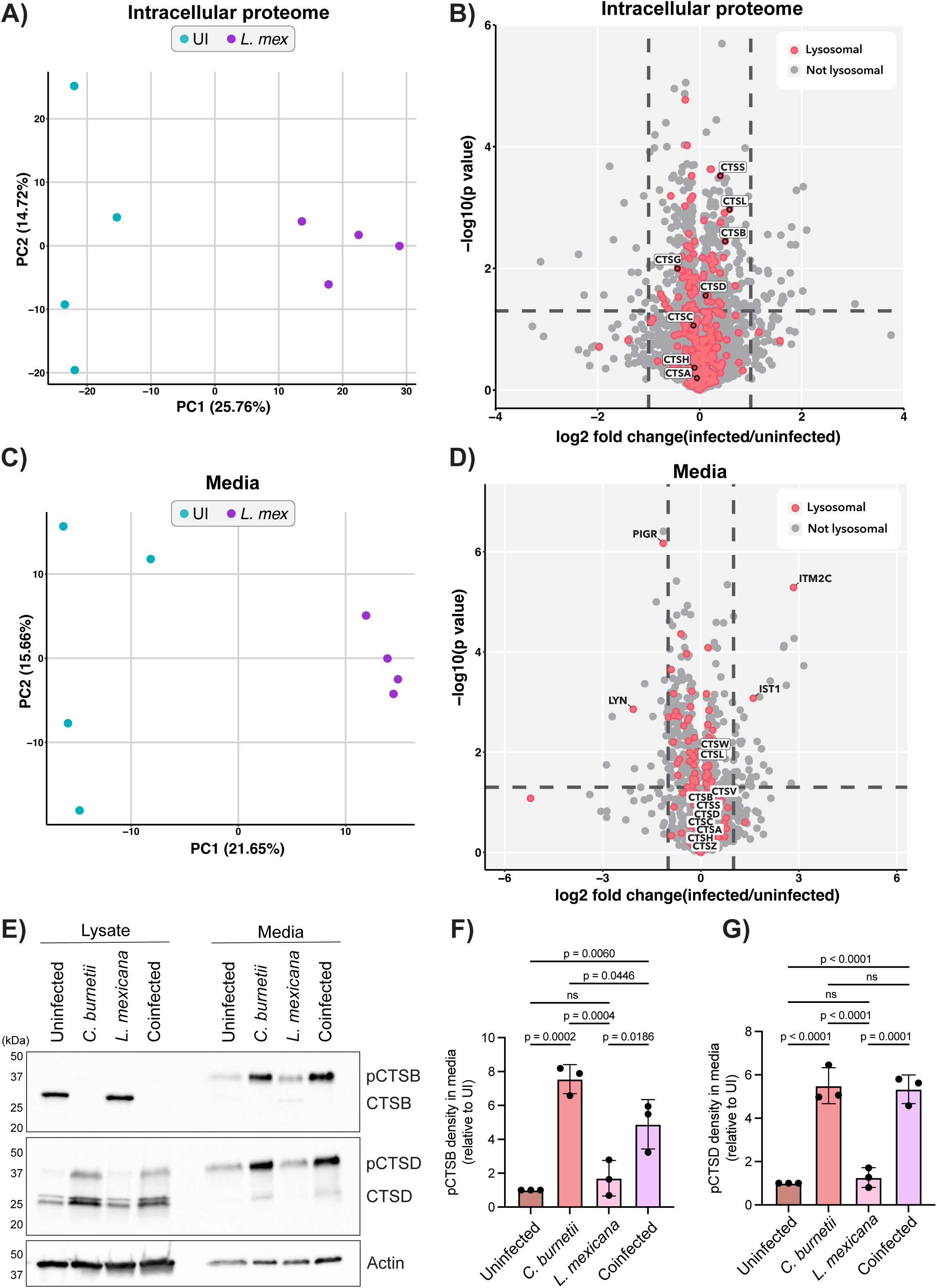
*L. mexicana* does not induce extracellular secretion of lysosomal proteins. **(A-D)** THP-1 cells were left uninfected (n=4) or infected with *L. mexicana* (n=4) at a MOI of 10 for 2 days before being washed and changed to serum-free culture media for a further 18 h. At this time, cell lysates and conditioned media were harvested and subjected to label-free quantitative mass spectrometry. **(A)** Principal component analysis (PCA) plot of proteomics data from whole cell lysates (“intracellular proteome”). Uninfected samples are shown in teal, *L. mexicana*-infected samples are shown in purple (n=4/group). **(B)** Volcano plot of proteins enriched in the intracellular proteome of *L. mexicana*-infected THP-1 cells as determined by proteomics analysis. *X*-axis represents log_2_(fold change) relative to uninfected cells, *Y*-axis shows statistical significance, -log_10_(p-value, as determined by student’s *t* test. Proteins with designated lysosomal localisation are coloured in pink, cathepsins and significantly altered lysosomal proteins are labelled with gene names. **(C)** As for (A) but depicting PCA plot of samples from conditioned media of *L. mexicana* infected THP-1 cells. **(D)** As for (B) but volcano plot of proteins enriched in the conditioned media from *L. mexicana*-infected THP-1 cells. **(E)** Immunoblot depicting cathepsin expression in the lysates and media of cells infected with *C. burnetii, L. mexicana* or co-infected with both microbes simultaneously. Blot is representative of 3 independent experiments, quantified in **(F, G). (F)** Quantification of pro-cathepsin B (pCTSB) density in media as presented in (E). Values are expressed relative to uninfected and standardised against respective actin expression. Graph depicts 3 biological replicates. Error bars denote SD. **(G)** As for (F) but depicting pro-cathepsin D (pCTSD) density in media. *P* values calculated using one-way ANOVA with Tukey’s correction. Mass spectrometry source data are available via ProteomeXchange with accession number PXD063745.

The proteomics analysis was extended to the conditioned media of *L. mexicana* infected cells to determine if lysosomal content was secreted to the extracellular space during infection, as observed for *C. burnetii*. 1,642 proteins were identified and quantified in the conditioned media fraction, 64 of which were significantly enriched in abundance during *L. mexicana* infection (as determined using a student’s t-test with *p*<0.05 and log_2_ fold change>1). As before, principal component analysis confirmed that biological replicates from each group clustered together (Fig 5C). There was no significant enrichment of lysosomal proteins in the media following *L. mexicana* infection (Fig 5D, pink circles). Further to this, cathepsins were broadly unchanged in abundance in the media from *L. mexicana-*infected cells when compared to media harvested from uninfected cells (Fig 5D, labels). Collectively, this demonstrates that secretion of lysosomal content is not observed upon infection with all intralysosomal pathogens.

Finally, cathepsin secretion following co-infection of cells with both *C. burnetii* and *L. mexicana* was examined. Immunoblotting revealed that, consistent with previous data, *C. burnetii* infection induced a significant increase in the secretion of cathepsins B and D relative to uninfected cells (Fig 5E, F, G). This increase was not observed during infection of cells with *L. mexicana* alone, aligning with the mass spectrometry results. Interestingly, when cells were coinfected with both pathogens at equal MOI, *C. burnetii* appeared to dominate with respect to cathepsin trafficking (Fig 5E, F, G). In cell lysates, cathepsin B was removed and immature cathepsin D (pCTSD) accumulated, while in conditioned media, both cathepsins were increased in abundance. Overall, the findings presented here imply that modulation of cathepsins is not shared between all intralysosomal pathogens. Instead, it is a specific virulence strategy induced by *C. burnetii*, while *L. mexicana* amastigotes are able to tolerate exposure to the full repertoire of host lysosomal proteases.

## DISCUSSION

The lysosome represents a key innate immune strategy for the removal of invading pathogens. Yet, a unique subset of intracellular pathogens can avoid degradation by these mechanisms and replicate in phagolysosomal compartments. We recently reported that the intralysosomal bacterium *Coxiella burnetii*, the causative agent of Q fever, strategically modifies the protease repertoire of the lysosome by removing the key enzyme cathepsin B to promote bacterial success [21]. In the present study, we examined whether this extended to *Leishmania*, a protozoan parasite that also resides within an intralysosomal niche.

We examined the abundance and activity of cathepsin proteases during *C. burnetii* or *L. mexicana* infection using immunoblotting and activity-based probe analysis. Loss of cathepsin B during *C. burnetii* infection was consistently observed, however this was not the case for cells infected with *L. mexicana* (Fig 1). In fact, the protease cohort of *L. mexicana-* infected cells appeared broadly unchanged from uninfected cells. This suggests that cathepsin B removal is a specific virulence strategy employed by *C. burnetii* to promote intracellular success. Although the vacuoles of *C. burnetii* and *L. mexicana* are alike in being fusogenic, acidic compartments, there is existing literature to suggest that these two pathogens exert differing effects on the host cell [28]. This research aligns with these observations, adding that this extends to modulation of the host lysosomal protease cohort. Furthermore, this examination of protease abundance and activity in *L. mexicana* infection is also consistent with previous reports using *L. amazonensis*, which showed that lysosomal hydrolase activity remains constant or is marginally elevated during infection of bone-marrow derived macrophages [12].

A question that remains to be addressed is whether the lysosomal environment is altered during *L. mexicana* infection. Early studies showed that surface lipophosphoglycan (LPG) of promastigote stages of *L. donovani* and *L. major* can delay phagosome maturation after entry into macrophages, allowing time for parasites to differentiate to the hydrolase-resistant amastigote form [29]. Phagosome maturation was delayed, in part, by exclusion of v-ATPase, allowing *Leishmania* promastigotes to establish infection in a non-acidified compartment [30]. However, other species of *Leishmania* (e.g. *L. mexicana*) are not dependent on LPG for promastigote infection and intracellular development, indicating species-specific mechanisms for avoiding or preventing lysosomal killing [32]. Moreover, in contrast to promastigotes, *Leishmania* amastigotes lack LPG [31] and there is no evidence that these stages induce delayed phagolysosome maturation in any species [9]. Based on the findings of this study we propose that neither promastigotes or amastigotes of *Leishmania* are dependent on the down-regulation of host cathepsin proteases for intracellular survival.

Here both co- and super-infection models were established to determine if the presence of both pathogens inside a cell might alter cathepsin B abundance or pathogen proliferation. In these models, co-infected cells appeared to display the same cathepsin phenotypes as cells infected with *C. burnetii* alone, suggesting that *C. burnetii* orchestrates changes to cathepsins that *L. mexicana* can withstand without any apparent fitness cost. Importantly, co-infection with *L. mexicana* did not rescue the loss of cathepsin B observed during *C. burnetii* infection. It should be acknowledged that the ability of these pathogens to replicate inside a THP-1 cell *in vitro* varies significantly. In existing literature, *C. burnetii* shows 500 to 1000-fold replication across 5 days of infection [33, 34], while THP-1 cells only support a much smaller number (∼20-fold replication) of *L. mexicana* parasites [35].

Although *C. burnetii* and *L. mexicana* show similarities in their intracellular niches, it is interesting to consider the key differences between these two pathogens. Specifically, *Leishmania* amastigotes switch to a slow growth state with greatly reduced rates of metabolic activity compared to extracellular promastigotes, which is associated with increased resistance to a wide variety of cellular and oxidative stresses (reviewed in [36, 37]). Interestingly, both pathogens are auxotrophic for many amino acids, and it will be of interest to determine whether alterations in proteolytic capacity within the phagolysosome are associated with changes in the availability of free amino acids/peptides. The lysosome-localised host mechanistic target of rapamycin complex 1 (mTORC1) is a key regulator of cellular nutrient sensing but also has a role in lysosomal homeostasis by controlling the activity of TFEB, the major regulator of lysosomal biogenesis and autophagy [38, 39]. Under nutrient-rich conditions, mTORC1 phosphorylates TFEB to keep it retained in the cytoplasm, maintaining homeostasis by preventing excess lysosomal biogenesis and autophagy. However, the *Leishmania* protease GP63 cleaves mTOR, preventing proper host protein translation and effectively reducing the host immune response [40]. Theoretically, mTOR cleavage by *Leishmania* should result in TFEB dephosphorylation and nuclear translocation, culminating in increased lysosomal biogenesis and autophagy. Although we did not directly examine TFEB activity in this study, the proteomics analysis performed in Fig 5 did not show increased abundance of lysosomal proteins during *L. mexicana* infection. It is therefore interesting to consider whether other regulatory pathways such as calcium/calcineurin signalling, JNK signalling or the AMP-activated protein kinase (AMPK) pathway contribute to *Leishmania* modulation of the host cell.

Collectively, alterations to lysosomal biology that are induced by *C. burnetii* are not observed during infection with *L. mexicana*. A potential rationale for this is that the unique metabolic state of *Leishmania* allows it to withstand the harsh phagolysosomal environment, while *C. burnetii* instead resorts to altering the environment itself through modulation of key lysosomal enzymes. Cumulatively, the findings presented here provides deeper insight into how these two “intralysosomal” pathogens differ in their mechanisms of withstanding this hostile environment.

## MATERIALS AND METHODS

### Propagation of C. burnetii and L. mexicana

Axenic cultures of *C. burnetii* Nine Mile Phase II RSA439 were generated in acidified cysteine citrate media 2 (ACCM2) [41, 42] for 6-7 days at 37 °C with 5% CO_2_ and 2.5% O_2_. Where required for cultivation of *C. burnetii icmL*::Tn, cultures were supplemented with kanamycin at 350 μg/ml.

Promastigote cultures of *L. mexicana* were grown in Roswell Park Memorial Institute (RPMI) media supplemented with 10% heat-inactivated fetal calf serum (FCS). Promastigotes were maintained at 27 °C and passaged every 3-4 days.

*L. mexicana* amastigotes were generated by transferring stationary phase promastigotes to RPMI-HCl (pH 5.5) with 20% FCS and incubating at 33 °C, 5% CO_2_ for 4 days prior to infection.

### Mammalian cell culture

THP-1 human monocytic cells were maintained in RPMI media with 10% FCS at 37 °C with 5% CO_2_. Cells were differentiated to macrophage-like cells with phorbol 12-myristate 13-acetate (PMA) at a concentration of 10^-8^ M for 72 h prior to infection.

### Immunoblotting

At the desired time-points post infection, cells were washed with PBS and harvested in 2X Laemmli buffer for SDS-PAGE. Samples were boiled at 95 °C for 10 min before being electrophoresed on an Any kD^TM^ Mini-PROTEAN® TGX Stain-Free^TM^ protein gel (Bio-Rad) at 165 V for 45 min. Gels were transferred to PVDF membranes and blocked for 1 h in 5% skim milk in Tris-buffered saline + 0.1% Tween-20 (TBST). Primary and secondary antibodies (see below) were diluted in 5% bovine serum albumin (BSA) in TBST. Blots were imaged on an AI680 Imager (GE Healthcare) or a ChemiDoc MP Imager (Bio-Rad) using the Western ECL Clarity substrate (Bio-Rad).

Antibodies used in this study are as follows: anti-Cathepsin B (Cell Signalling Technology, #31718, 1:2000), anti-Cathepsin C (Santa Cruz Biotechnology, #sc-74590, 1:2000, anti-Cathepsin D (Abcam, #72915, 1:5000), Actin (Sigma, #A1978, 1:10,000).

### Activity-based probe analysis

At the desired time-point, cells were harvested by replacing media with PBS and using a cell scraper to collect. Cells were pelleted by centrifugation at 7000 x g for 5 min. Cells were lysed in a citrate buffer (50 mM citrate, pH 5.5, 0.5% CHAPS, 0.1% Triton-X 100, 4 mM DTT) and total protein was quantified using a BCA assay (Pierce). BMV109 (cathepsins B, S, X, L) was added to lysates (1 µM) and incubated for 45 min at 37 °C. The reaction was stopped by addition of sample buffer and boiling for 5 min. Proteins were resolved via SDS-PAGE and in-gel fluorescence was detected by scanning on a Typhoon imager (GE Healthcare) using a Cy5 filter. When required, gels were then transferred to nitrocellulose membranes and immunoblotted as above.

### Infection of THP-1 cells with *L. mexicana* and/or *C. burnetii*

THP-1 cells were plated at a density of 5 x 10^5^ cells per well in a 24-well plate and differentiated with PMA as above.

For *C. burnetii* infections, stationary phase (6 or 7-day) axenic cultures were pelleted and resuspended in RPMI + 10% FCS before being quantitated using qPCR and primers specific to *ompA*. Concentration of the bacterial culture was determined by comparison against a standard curve. *C. burnetii* was added to THP-1 cells at MOI 10 before culture plates were centrifuged for 30 min at 500 x g. Cells were then washed thrice with PBS and incubated at 37 °C, 5% CO_2_ until the desired time-point.

For *L. mexicana* infections, amastigote cultures were quantified visually using a Neubauer haemocytometer. Parasite suspensions were added to cells at a MOI of 5 or 10 before plates were centrifuged at 300 x g for 5 min to promote contact between amastigotes and THP-1 cells. Cells were left to incubate at 33 °C with 5% CO_2_ for 4 hrs before being washed thrice with PBS and returned to 33 °C, 5% CO_2_ for the desired time.

For co-infections, bacteria and parasites were individually quantified as above. THP-1 cells were either infected with both organisms simultaneously (co-infection) or infected with *L. mexicana* and left for 3 d at 33 °C before being infected with *C. burnetii* (super-infection).

### Quantification of intracellular replication

Quantitative PCR was used to determine intracellular replication of *C. burnetii* and *L. mexicana* within THP-1 cells. Immediately following infection (D0) or at days 1, 3, and 5 post infection, cells were osmotically lysed in dH_2_O and collected by centrifugation at 17,000 x g for 15 min. Genomic DNA was extracted using Quick DNA miniprep kit (Zymo Research) according to the manufacturer’s protocol. qPCR was performed using the SensiFast SYBR No-Rox kit (Bioline) and primers specific to either *ompA* (for *C. burnetii*) or *its1* (for *L. mexicana*). Serial dilutions of *C. burnetii* or *L. mexicana* were used to generate a standard curve for quantitation. qPCR was performed using a QuantStudio 7 Flex RT-qPCR (Thermo Fisher Scientific).

Oligonucleotides used for quantification are as follows: OmpA_F CAGAGCCGGGAGTCAAGCT, OmpA_R CTGAGTAGGAGATTTGAATCGC, Its1_F GGATCATTTTCCGATGATTACACC [26], Its1_R CTGCAAATGTTGTTTTTGAGTACA [26].

### Mass spectrometry on *L. mexicana* infected cell lysates and culture media Sample collection

THP-1 cells were plated in 10 cm dishes at a density of 1.1 x 10^7^ cells per dish and infected with *L. mexicana* at a MOI of 25, as described above. 4x biological replicates were prepared for each condition (uninfected and infected, respectively). After 48 h of infection, cells were washed once with PBS and incubated in fresh RPMI containing no FCS (serum-free) and incubated for a further 18 h. After this time, supernatants were collected and centrifuged at 3000 x g for 10 min to pellet any cell debris. The supernatants were then concentrated through Amicon® Ultra Centrifugal Filters (Merck) with a molecular weight cut off of 10 kDa, before being combined with a lysis buffer containing 10% sodium dodecyl sulfate (SDS) and 100 mM Tris-HCl, pH 8.0. Simultaneously, cell fractions were washed thrice with PBS before being lysed in the same lysis buffer. All samples were then incubated at 95 °C with shaking at 2000 rpm to shear DNA before total protein was quantitated using Nanodrop. Mass spectrometry sample preparation was performed using 100 μg of protein as input.

### Sample preparation

Dithiothreitol (DTT) was added to a final concentration of 10 mM and samples were incubated at 95 °C for 10 min before alkylation was performed using 40 mM iodoacetamide at room temperature in the dark. After 30 min, the reaction was quenched with the addition of another 10 mM DTT, then samples were acidified with 12% phosphoric acid. Samples were then combined with a binding/wash buffer (100 mM Tris (pH 8.0), 90% (v/v) methanol), before being loaded onto S-trap micro columns (ProtiFi, USA). Columns were washed four times with the same binding/wash buffer before samples were digested with SoluTrypsin (Sigma) using a 3 µg trypsin per sample (approximately 1:33 trypsin:input), diluted in 50 mM Tris-HCl pH 8. Digestion was performed at 37 °C in a humidified container for 18 h. The following day, samples were eluted by centrifugation using 50 mM Tris-HCl pH 8, followed by 0.2% formic acid, then with 50% acetonitrile and 0.2% formic acid at 4000 x g for 1 min each. The combined eluate was collected and vacuum dried before being resuspended in mass spectrometry loading buffer (“Buffer A*”, 0.1% trifluoroacetic acid, 2% acetonitrile) and subjected to desalting using C_18_ stage tips [43, 44].

### Liquid chromatography mass spectrometry

C_18_-enriched proteome samples were re-suspended in Buffer A* and separated using a two-column chromatography setup composed of a PepMap100 C_18_ 20-mm by 75-mm trap (Thermo Fisher Scientific) and a PepMap C_18_ 500-mm by 75-mm analytical column (Thermo Fisher Scientific) using a Dionex Ultimate 3000 UPLC (Thermo Fisher Scientific). Samples were concentrated onto the trap column at 5 µL/min for 6 min with Buffer A (0.1% formic acid, 2% DMSO) and then infused into an Orbitrap Fusion Lumos™ (Thermo Fisher Scientific) at 300 nl/mi via the analytical columns. Peptides were separated by altering the buffer composition from 3% Buffer B (0.1% formic acid, 77.9% acetonitrile, 2% DMSO) to 23% B over 70 min, then from 23% B to 40% B over 4 min and then from 40% B to 80% B over 3 min. The composition was held at 80% B for 2 min before being returned to 3% B for 10 min. The Orbitrap Fusion Lumos™ Mass Spectrometer was operated in a data-dependent mode automatically switching between the acquisition of a single Orbitrap MS scan (300-2000 m/z, maximal injection time of 50 ms, an Automated Gain Control (AGC) set to a maximum of 400k and a resolution of 60k) and 3 s of Orbitrap MS/MS HCD scans of precursors (NCE 35%, a maximal injection time of 80 ms, a AGC of 125000 and a resolution of 30k).

### Data analysis

Raw spectral files were searched using FragPipe [45, 46] (v22.0) against *Homo sapiens* (UniProt accession: UP000005640) and *Leishmania mexicana* (UniProt accession: UP000007259) reference proteomes. FragPipe parameters were left on default unless otherwise specified. Label-free quantification (LFQ) was performed allowing for cysteine carbamidomethylation as a fixed modification (+57.0215 Da) and N-terminal acetylation (+42.0106 Da) and oxidation of methionine (+15.9949 Da) as variable modifications. Protease specificity was ‘stricttrypsin’ with cleavage at K or R residues, and 2 missed cleavages (maximum) were allowed. Philosopher was used to calculate protein and peptide false discovery rates (FDRs). Resulting data was further analysed using the Perseus[47] (v.1.6.0.7) software. Data were subject to log_2_ transformation and then filtered to only retain proteins with valid values identified in at least three out of four biological replicates in at least one group (i.e. uninfected/infected). Missing values were imputed based off a downshifted normal distribution (width=0.3σ, downshift = -1.8σ). Statistical comparison between groups was performed using a student’s *t* test and multiple hypothesis correction based on permutation-based FDR. Data visualisation was performed using the ggplot2 package in R (v.4.2.1).

## ACKNOWLEDGEMENTS

We wish to acknowledge the Mass Spectrometry and Proteomics Facility (MSPF) at the Bio21 Institute at the University of Melbourne for access to and maintenance of key resources. L.E.B., B.X., and E.N.S.M were supported by an RTP scholarship from the Australian Government. MJM was supported by an Australian National Health and Medical Research Council (NHMRC) Principal Research Fellowship. L.E.E-M. was funded by a grant from the Australian National Health and Medical Research Council (GNT2011119). Research in H.J.N.’s laboratory was funded by the Australian National Health and Medical Research Council (GNT2010841).

## DATA AVAILABILITY

Mass spectrometry raw data files have been uploaded to the ProteomeXchange consortium via the PRIDE [48] repository with the dataset identifiers PXD063745.

## REFERENCES

1. Vaughn, B. and Y. Abu Kwaik, Idiosyncratic Biogenesis of Intracellular Pathogens-Containing Vacuoles. Front Cell Infect Microbiol, 2021. 11: p. 722433.

2. Ray, K., et al., Life on the inside: the intracellular lifestyle of cytosolic bacteria. Nature Reviews Microbiology, 2009. 7(5): p. 333–340.

3. Torres-Guerrero, E., et al., Leishmaniasis: a review. F1000Res, 2017. 6: p. 750.

4. Burza, S., S.L. Croft, and M. Boelaert, Leishmaniasis. Lancet, 2018. 392(10151): p. 951–970.

5. Alemayehu, B. and M. Alemayehu, Leishmaniasis: a review on parasite, vector and reservoir host. Health Science Journal, 2017. 11(4): p. 1.

6. Peters, N.C., et al., In vivo imaging reveals an essential role for neutrophils in leishmaniasis transmitted by sand flies. Science, 2008. 321(5891): p. 970–974.

7. Van Zandbergen, G., et al., Cutting edge: neutrophil granulocyte serves as a vector for Leishmania entry into macrophages. The Journal of Immunology, 2004. 173(11): p. 6521–6525.

8. Real, F. and R.A. Mortara, The Diverse and Dynamic Nature of Leishmania Parasitophorous Vacuoles Studied by Multidimensional Imaging. PLOS Neglected Tropical Diseases, 2012. 6(2): p. e1518.

9. Courret, N., et al., Biogenesis of Leishmania-harbouring parasitophorous vacuoles following phagocytosis of the metacyclic promastigote or amastigote stages of the parasites. J Cell Sci, 2002. 115(Pt 11): p. 2303–16.

10. Lippuner, C., et al., Real-time imaging of Leishmania mexicana-infected early phagosomes: a study using primary macrophages generated from green fluorescent protein-Rab5 transgenic mice. Faseb j, 2009. 23(2): p. 483–91.

11. Antoine, J.C., et al., Leishmania mexicana: a cytochemical and quantitative study of lysosomal enzymes in infected rat bone marrow-derived macrophages. Exp Parasitol, 1987. 64(3): p. 485–98.

12. Prina, E., et al., Localization and activity of various lysosomal proteases in Leishmania amazonensis-infected macrophages. Infect Immun, 1990. 58(6): p. 1730–7.

13. Moradin, N. and A. Descoteaux, Leishmania promastigotes: building a safe niche within macrophages. Front Cell Infect Microbiol, 2012. 2: p. 121.

14. Hartzell, J.D., et al. Q fever: epidemiology, diagnosis, and treatment. in Mayo Clinic Proceedings. 2008. Elsevier.

15. Kohler, L.J. and C.R. Roy, Biogenesis of the lysosome-derived vacuole containing Coxiella burnetii. Microbes and Infection, 2015. 17(11): p. 766–771.

16. Larson, C.L., et al., Right on Q: genetics begin to unravel Coxiella burnetii host cell interactions. Future Microbiol, 2016. 11(7): p. 919–39.

17. Andreoli, W.K. and R.A. Mortara, Acidification modulates the traffic of Trypanosoma cruzi trypomastigotes in Vero cells harbouring Coxiella burnetii vacuoles. International Journal for Parasitology, 2003. 33(2): p. 185–197.

18. de Chastellier, C., M. Thibon, and M. Rabinovitch, Construction of chimeric phagosomes that shelter Mycobacterium avium and Coxiella burnetii (phase II) in doubly infected mouse macrophages: an ultrastructural study. Eur J Cell Biol, 1999. 78(8): p. 580–92.

19. Veras, P.S., et al., Entry and survival of Leishmania amazonensis amastigotes within phagolysosome-like vacuoles that shelter Coxiella burnetii in Chinese hamster ovary cells. Infection and Immunity, 1995. 63(9): p. 3502–3506.

20. Beare, P.A., et al., Dot/Icm Type IVB Secretion System Requirements for Coxiella burnetii Growth in Human Macrophages. mBio, 2011. 2(4): p. 10.1128/mbio.00175-11.

21. Bird, L.E., et al., Coxiella burnetii manipulates the lysosomal protease cathepsin B to facilitate intracellular success. Nature Communications, 2025. 16(1): p. 3844.

22. Edgington-Mitchell, L.E., M. Bogyo, and M. Verdoes, Live cell imaging and profiling of cysteine cathepsin activity using a quenched activity-based probe, in Activity-Based Proteomics: Methods and Protocols, H.S. Overkleeft and B.I. Florea, Editors. 2017, Springer New York: New York, NY. p. 145–159.

23. Verdoes, M., et al., Improved quenched fluorescent probe for imaging of cysteine cathepsin activity. J Am Chem Soc, 2013. 135(39): p. 14726–30.

24. Sanman, L.E. and M. Bogyo, Activity-Based Profiling of Proteases. Annual Review of Biochemistry, 2014. 83(Volume 83, 2014): p. 249–273.

25. Lau, N., et al., Perturbation of ATG16L1 function impairs the biogenesis of Salmonella and Coxiella replication vacuoles. Mol Microbiol, 2022. 117(2): p. 235–251.

26. Ceccarelli, M., et al., Differentiation of Leishmania (L.) infantum, Leishmania (L.) amazonensis and Leishmania (L.) mexicana Using Sequential qPCR Assays and High-Resolution Melt Analysis. Microorganisms, 2020. 8(6).

27. Padmanabhan, B., et al., Biogenesis of the spacious Coxiella-containing vacuole depends on host transcription factors TFEB and TFE3. Infect Immun, 2020. 88(3): p. e00534–19.

28. Millar, J.A., et al., Coxiella burnetii and Leishmania mexicana residing within similar parasitophorous vacuoles elicit disparate host responses. Front Microbiol, 2015. 6: p. 794.

29. Desjardins, M. and A. Descoteaux, Inhibition of phagolysosomal biogenesis by the Leishmania lipophosphoglycan. Journal of Experimental Medicine, 1997. 185(12): p. 2061–2068.

30. Vinet, A.F., et al., The Leishmania donovani Lipophosphoglycan Excludes the Vesicular Proton-ATPase from Phagosomes by Impairing the Recruitment of Synaptotagmin V. PLOS Pathogens, 2009. 5(10): p. e1000628.

31. Mcconville, M.J. and J.M. Blackwell, Developmental changes in the glycosylated phosphatidylinositols of Leishmania donovani. Characterization of the promastigote and amastigote glycolipids. Journal of Biological Chemistry, 1991. 266(23): p. 15170–15179.

32. Ilg, T., Lipophosphoglycan is not required for infection of macrophages or mice by Leishmania mexicana. The EMBO journal, 2000.

33. Loterio, R.K., et al., Coxiella co-opts the Glutathione Peroxidase 4 to protect the host cell from oxidative stress-induced cell death. Proc Natl Acad Sci U S A, 2023. 120(36): p. e2308752120.

34. Thomas, D.R., et al., Coxiella burnetii protein CBU2016 supports CCV expansion. Pathogens and Disease, 2024. 82: p. ftae018.

35. Dagley, M.J., et al., High-Content Assay for Measuring Intracellular Growth of Leishmania in Human Macrophages. ASSAY and Drug Development Technologies, 2015. 13(7): p. 389–401.

36. McConville, M.J., et al., Leishmania carbon metabolism in the macrophage phagolysosome-feast or famine? F1000Res, 2015. 4(F1000 Faculty Rev): p. 938.

37. Saunders, E.C., et al., Metabolic stringent response in intracellular stages of Leishmania. Current Opinion in Microbiology, 2021. 63: p. 126–132.

38. Martina, J.A., et al., MTORC1 functions as a transcriptional regulator of autophagy by preventing nuclear transport of TFEB. Autophagy, 2012. 8(6): p. 903–14.

39. Palmieri, M., et al., Characterization of the CLEAR network reveals an integrated control of cellular clearance pathways. Hum Mol Genet, 2011. 20(19): p. 3852–66.

40. Jaramillo, M., et al., Leishmania repression of host translation through mTOR cleavage is required for parasite survival and infection. Cell Host Microbe, 2011. 9(4): p. 331–41.

41. Omsland, A., et al., Isolation from animal tissue and genetic transformation of Coxiella burnetii are facilitated by an improved axenic growth medium. Appl Environ Microbiol, 2011. 77(11): p. 3720–5.

42. Omsland, A., et al., Host cell-free growth of the Q fever bacterium Coxiella burnetii. Proceedings of the National Academy of Sciences, 2009. 106(11): p. 4430–4434.

43. Rappsilber, J., Y. Ishihama, and M. Mann, Stop and go extraction tips for matrix-assisted laser desorption/ionization, nanoelectrospray, and LC/MS sample pretreatment in proteomics. Anal Chem, 2003. 75(3): p. 663–70.

44. Rappsilber, J., M. Mann, and Y. Ishihama, Protocol for micro-purification, enrichment, pre-fractionation and storage of peptides for proteomics using StageTips. Nat Protoc, 2007. 2(8): p. 1896–906.

45. Kong, A.T., et al., MSFragger: ultrafast and comprehensive peptide identification in mass spectrometry–based proteomics. Nature Methods, 2017. 14(5): p. 513–520.

46. Yu, F., S.E. Haynes, and A.I. Nesvizhskii, IonQuant Enables Accurate and Sensitive Label-Free Quantification With FDR-Controlled Match-Between-Runs. Molecular & Cellular Proteomics, 2021. 20.

47. Tyanova, S., et al., The Perseus computational platform for comprehensive analysis of (prote)omics data. Nature Methods, 2016. 13(9): p. 731–740.

48. Perez-Riverol, Y., et al., The PRIDE database resources in 2022: A hub for mass spectrometry-based proteomics evidences. Nucleic Acids Research, 2022. 50(D1): p. D543–D552.

